# Deep Brain Ultrasound Augments Human Attention

**DOI:** 10.1101/2025.09.28.678996

**Authors:** Hamidreza Ramezanpour, Ghazaleh Darmani, Regina Annirood, Can Sarica, Jean-Francois Nankoo, Samuel Pichardo, Jeffrey Schall, Andres M. Lozano, Robert Chen

## Abstract

**Background:** Deep brain ultrasound offers a novel means of modulating human cognition by noninvasively targeting subcortical structures that were previously accessible only through invasive procedures. While decades of research have mapped cortical circuits of attention, the causal roles of deep hubs such as the basal ganglia and thalamus remain poorly understood in the healthy human brain.

**Objectives/Hypothesis:** To test whether low intensity transcranial ultrasound stimulation (TUS) of two nodes in the basal ganglia-thalamic network, the globus pallidus internus (GPi) and the pulvinar, causally alters visual attention. We hypothesized that TUS-induced modulations in attentional performance would be site specific, reflecting distinct circuit functions.

**Results:** Across sessions, focal TUS accelerated reaction time in a visual search task, indicating augmented attention. Reaction time improvements were observed after stimulation relative to baseline. A dissociation emerged across sites: both GPi and pulvinar enhanced reaction times, but pulvinar yielded more robust benefits for target present trials at peripheral eccentricities, and improved search efficiency in the same trials.

**Conclusions:** These findings provide causal evidence that human attentional control can be steered at deep subcortical sites. TUS offers a practical approach for dissecting circuit level contributions to cognition and a potential noninvasive avenue for enhancing attention and other cognitive or affective functions.

## 1. Introduction

Decades of research have mapped the cortical networks underlying attention in humans[1–3]. However, most human studies of attention have been correlational or based on neurological patients with lesions[4–8], which limits causal inference about underlying mechanisms. Deep subcortical structures like the basal ganglia and thalamus have been difficult to probe in healthy individuals because they lie beyond the reach of conventional non-invasive methods. As a result, the contributions of these deep hubs to attention remain poorly understood in the intact human brain. Developing methods to causally and focally modulate deep brain activity in humans would fill a critical gap between invasive patient studies and surface-focused neurostimulation, enabling new insights into how subcortical circuits actively shape cognition.

Transcranial ultrasound stimulation (TUS) has recently emerged as a technology uniquely suited for non-invasive, focal perturbation of deep brain circuits[9]. Appropriate TUS protocols can induce neural effects that outlast the stimulation period by tens of minutes, permitting the study of “offline” changes in behavior[10,11]. These features position TUS as a means to causally test the roles of deep brain structures in cognition for the first time in neurologically normal humans. Early applications in animals and humans have validated that TUS can modulate activity in subcortical targets such as the basal ganglia and thalamus[12,13], supporting its potential to serve as a non-invasive analog of deep brain stimulation (DBS).

The thalamus, long regarded as a passive relay for sensory information, is increasingly recognized as an active participant in cognitive processes[14]. Thalamic nuclei have extensive reciprocal connections with cortex, placing them in a strategic position to regulate information flow across distributed networks[15]. In particular, the pulvinar nucleus of the thalamus has been implicated in visual attention[6,8,16]. The pulvinar is well-situated to influence attentional orienting and filtering of visual input, given its dense interconnections with visual cortical areas[17]. Consistent with this anatomy, a growing body of work in both non-human primates and humans links pulvinar activity to selective attention[18–21]. For example, neural recordings and perturbations in monkeys have shown that pulvinar synchronizes cortical activity and can bias visual processing during attention tasks[22]. Inactivation of the pulvinar in monkeys impairs target detection[23], whereas driving pulvinar activity can enhance attention and performance[24]. Together, these studies suggest that the thalamus – and the pulvinar in particular – plays a key role in coordinating spatial attention, rather than serving as a mere relay.

By contrast, the globus pallidus internus (GPi), the main basal ganglia output nucleus, has classically been associated with motor functions[25] (for instance, GPi is a primary surgical target for movement disorders[26,27]). Its potential contributions to cognition have received comparatively limited attention. Anatomical and circuit-tracing studies demonstrate that the basal ganglia are not exclusively motor: they send projections to the prefrontal cortex and other non-motor regions[28]. Indeed, the primate GPi is a node in fronto-striatal loops that could influence decision-making and cognitive control[29,30]. Neurophysiological observations in clinical contexts also hint at a cognitive role for GPi – for example, GPi neurons modulate their activity during oddball attention tasks in Parkinson’s patients[31]. Yet, direct evidence for GPi involvement in attentional processes has remained elusive, in part because non-invasive methods to selectively perturb the GPi have not been available. The advent of TUS now enables tests of whether this deep motor structure also contributes to higher-order cognitive operations.

Against this backdrop, we leveraged low-intensity TUS to causally modulate the pulvinar and the GPi in healthy human participants performing a visual search. By targeting these subcortical structures, our aim was to determine how stimulating thalamic versus basal ganglia nodes alters top-down attention.

## 2. Materials and Methods

### 2.1. Participants

Fifteen healthy adult volunteers (age 18–35 years; 4 male, 11 female) participated in the study. All were neurologically healthy, had normal or corrected-to-normal vision, and provided written informed consent. The University Health Network Research Ethics Board approved the study in accordance with the Declaration of Helsinki. The same cohort of participants had previously taken part in our study on GPi and pulvinar stimulation during a stop-signal task[12], enabling us to use identical stimulation and targeting procedures here.

### 2.2. Ultrasound Stimulation and Acoustic Modelling

Stimulation was delivered with a single-element, 500 kHz focused transducer (Sonic Concepts, Bothell, WA) coupled to the scalp via ultrasound gel. Each session targeted one structure bilaterally, the internal segment of the globus pallidus (GPi) or the pulvinar nucleus of the thalamus, using neuronavigation (Brainsight, Rogue Research, Montreal, Canada). The order was randomized across participants and two sessions conducted at least one week apart. A theta-burst transcranial ultrasound stimulation (tbTUS) protocol was applied, consisting of 20 ms bursts at 50 Hz repeated every 200 ms (10% duty cycle) for 120 s per hemisphere. The spatial peak pulse average intensity in water was 30 W/cm^2^.

Target locations were identified from individual T1- and T2-weighted MRI scans, co-registered to a standard atlas (AAL3) and refined to each participant’s anatomy. For GPi, average MNI coordinates were ±20, −8, −4; for pulvinar, ±11, −33, 6. Coordinates were transformed into subject space and optimized for skull geometry using individualized acoustic simulations in BabelBrain[35]. Finite-difference time-domain methods incorporating skull density and cortical surfaces were used to estimate in-brain intensities and focal profiles. Across participants, modeled focal spots were ∼24 mm axially and 4– 5 mm laterally at −6 dB, with estimated in-brain ISPPA between 2–8 W/cm.

Acoustic simulations predicted minimal thermal effects, with maximum rises of 0.16 °C in the brain, 0.25 °C in the skin, and 0.30 °C in the skull. Transducer adjustments (Δx, Δy, Δz) were required to compensate for skull-induced distortions, reducing the distance between intended and effective targets. Within the target region, ISPPA values ranged from 2.5–7 W/cm^2^ and ISPTA from 0.25–0.7 W/cm^2^, with mechanical indices between 0.38–0.67. All parameters remained well within FDA and ITRUSST safety limits[36].

### 2.3. Visual Search Task

The behavioral paradigm was a classic “T among Ls” visual search task. On each trial, a rotated “T” target was either present or absent among distractors. Set size was varied randomly across 1, 4, 8, 16, or 32 items. Target positions were randomized across eccentricities: central (<8°), intermediate (8–30°), and peripheral (>30°). Each trial began with a small central fixation cross that participants were instructed to hold for 500 ms, after which the target and distractors were displayed. Participants sat approximately 70 cm from the monitor and indicated, as quickly and accurately as possible, whether the target was present or absent using two different response keys on a keyboard. Trials were separated by a 1-s inter-trial interval, and responses were accepted within a maximum of 10 s. Each experimental session consisted of three blocks: Pre (baseline), Post1 (immediately following sonication of the first hemisphere), and Post2 (following sonication of the contralateral hemisphere). Each block contained ∼300 trials, balanced for target presence/absence and eccentricity. Although tbTUS effects typically emerge within 5 minutes and peak around 10–15 minutes post-exposure, the length of each block ensured overlap with this window. In addition, the second post-sonication block was conducted immediately after a second round of tbTUS, ensuring that stimulation effects were still present and not washed out, consistent with prior studies showing effects lasting up to one hour post-stimulation[10–12].

### 2.4. Data Analysis

Analyses were restricted to correct trials. Reaction times (RTs) were characterized with empirical cumulative distribution functions (ECDFs) at each set size. Pre vs Post1, Pre vs Post2, and Post1 vs Post2 comparisons were performed using two-sample Kolmogorov–Smirnov (KS) tests. Distributional shifts were further quantified as differences in median RT (ΔMedian).

Eccentricity effects were analyzed by computing ΔRT (Post–Pre) separately for central, intermediate, and peripheral bins. Within-site effects were tested against zero (one-sample t-tests) and between-site differences were assessed with paired-sample t-tests.

Search efficiency was estimated by regressing RTs against set size. The intercept reflected general response speed independent of load, while the slope indexed efficiency (s/item). ΔSlopes were tested within site and contrasted between sites. All p-values were corrected for multiple comparisons using the Benjamini–Hochberg false discovery rate (FDR, q = 0.05).

## 3. Results

Each participant completed a visual search task (Fig. 1A) in three blocks: baseline (Pre), shortly after the first stimulation (Post-1), and shortly after the second stimulation on the same day (Post-2), with TUS targeting either the GPi or the pulvinar bilaterally, with the order randomized across participants, in two sessions conducted at least one week apart. We applied a theta-burst TUS protocol, which has been shown to produce long-lasting effects of up to 1 hour after stimulation cessation. To ensure that spatial distribution of visual stimuli does not influence potential impacts of TUS across session, for each subject (n = 15), a 2D histogram of stimulus positions was computed separately for trials with targets (Fig. 1B) and distractors (Fig. 1C) during each session (pre, post1, post2). Gaze-reported or click-reported screen coordinates were binned into a 10 × 10 grid, using consistent bin edges derived from the full dataset. Each subject’s heatmap was normalized to its maximum bin value to account for individual differences in the number and distribution of trials. Group-level maps represent the average of normalized heatmaps across subjects. Warmer colors indicate bins with higher relative frequency. To test for session-related changes in spatial deployment, paired t-tests were performed at each bin between sessions, corrected using false discovery rate (FDR) at α = 0.05. No significant differences were observed across any pairwise session comparison for either targets or distractors, indicating that the spatial distribution of visual stimuli remained consistent across timepoints.

**Figure 1.**
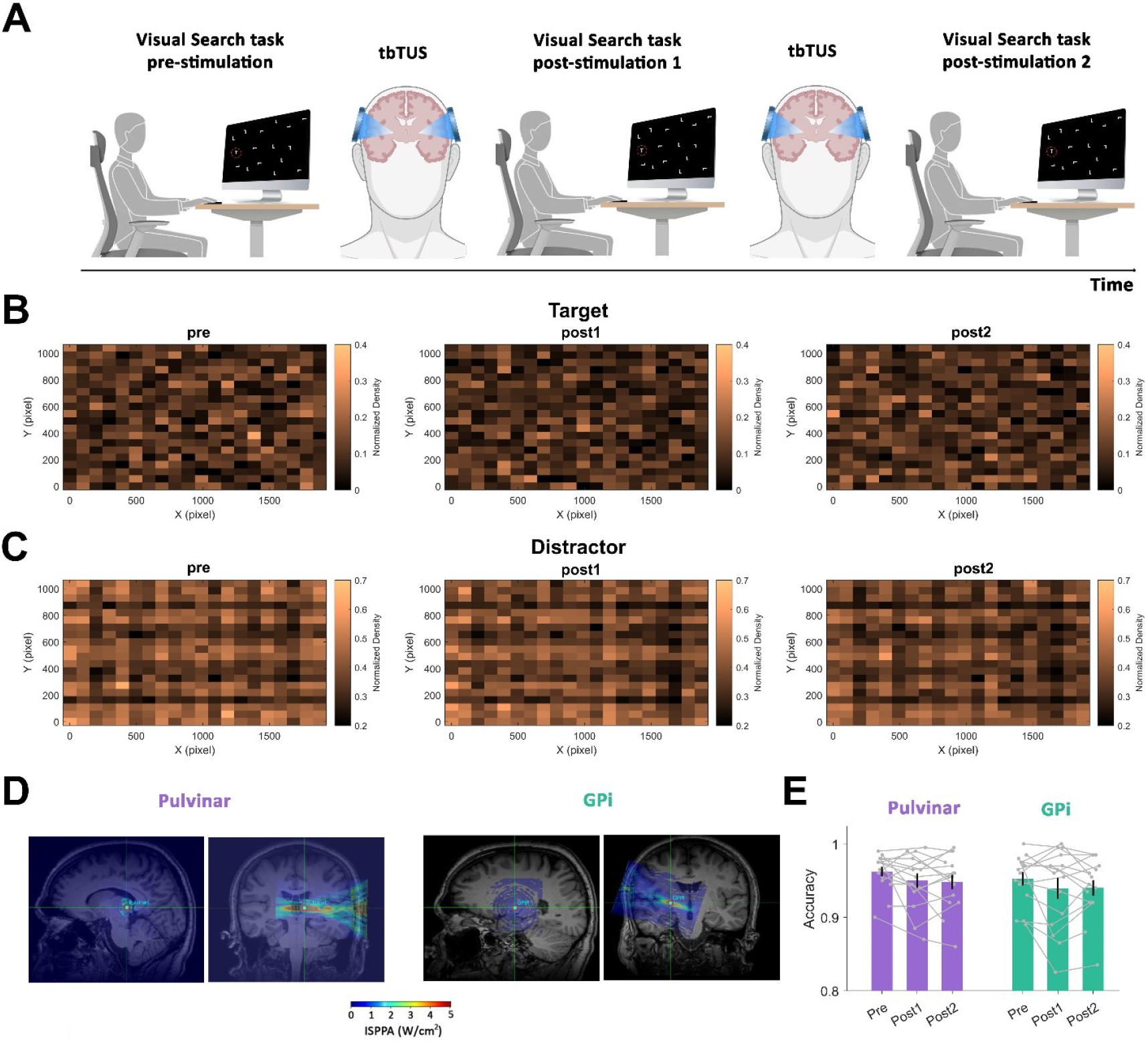
Effects of GPi and pulvinar TUS on visual search performance. (A) Experimental paradigm. Participants (n = 15) performed a “T among Ls” visual search task before and after bilateral theta-burst TUS (tbTUS) targeting either GPi or pulvinar. Each session included a baseline block (Pre), a block immediately after the first sonication (Post1), and a block after the second sonication (Post2). Set size (1, 4, 8, 16, or 32 items) and target eccentricity (<8°, 8–30°, >30°) were manipulated. (B-C) Group-level spatial heatmaps of target and distractor positions across sessions. No significant differences were observed across any pairwise session comparison for either targets or distractors, indicating that the spatial distribution of visual stimuli remained consistent across timepoints. (D) Individualized acoustic simulations confirmed focal sonication of GPi and pulvinar with at target intensities in the low-intensity neuromodulatory range (color scale = ISPPA, W/cm^2^). Across subjects, modeled ISPPA was 2–8 W/cm^2^. (E) Target detection accuracy (mean ± SEM) for pulvinar (purple) and GPi (green) remained high across sessions (GPi: Pre = 0.962 ± 0.026, Post1 = 0.950 ± 0.037, Post2 = 0.948 ± 0.038; Pulvinar: Pre = 0.952 ± 0.035, Post1 = 0.939 ± 0.054, Post2 = 0.940 ± 0.041). No within- or between-site differences were significant after FDR correction (all p > 0.5). Bars and shaded error margins indicate SEM.

Individualized acoustic simulations, confirming focal sonication of the intended deep target with in-brain intensities in the safe low-intensity neuromodulatory range, are depicted in Fig. 1D. More detailed descriptions of the acoustic simulation outcomes for individual subjects are provided in our previous work^4^, in which the same cohort of participants underwent sonication of the same targets, with cognitive performance assessed before and after stimulation using a different paradigm (see also Methods for additional details on the task, TUS protocol, acoustic modeling, safety, and data analysis).

Task accuracy remained high across all sessions and was not affected by stimulation (Fig. 1E), indicating that TUS did not affect the probability of finding the target. Reaction time (RT) analyses, however, revealed a richer pattern.

Reaction time distributions were reliably accelerated by stimulation (Fig. 2A–D). This is most directly visible in the raw cumulative distributions (Fig. 3A for GPi and Fig. 3B for pulvinar), where the Post1 and Post2 curves are systematically shifted leftward relative to baseline, indicating faster responses. In Fig. 2A–D, we plot the subtraction of these curves (Post – Pre), which makes the size and timing of the differences clearer. Notably, these curves diverged most strongly at higher set sizes, consistent with greater modulation under increased task demands. Importantly, a similar pattern was observed when target-present and target-absent trials were analyzed separately, as shown in Fig.3. To exclude the possibility that the observed RT improvements reflected learning or task familiarity, we compared average RTs from the first and second halves of trials within each block. Across all blocks, in both GPi and pulvinar sessions and at both pre and post stimulation, no significant differences were found between early and late trials (all p > 0.05, Wilcoxon signed-rank test). This indicates that practice effects did not drive the speeding of responses, supporting the conclusion that TUS was responsible for the shortened reaction times.

**Figure 2.**
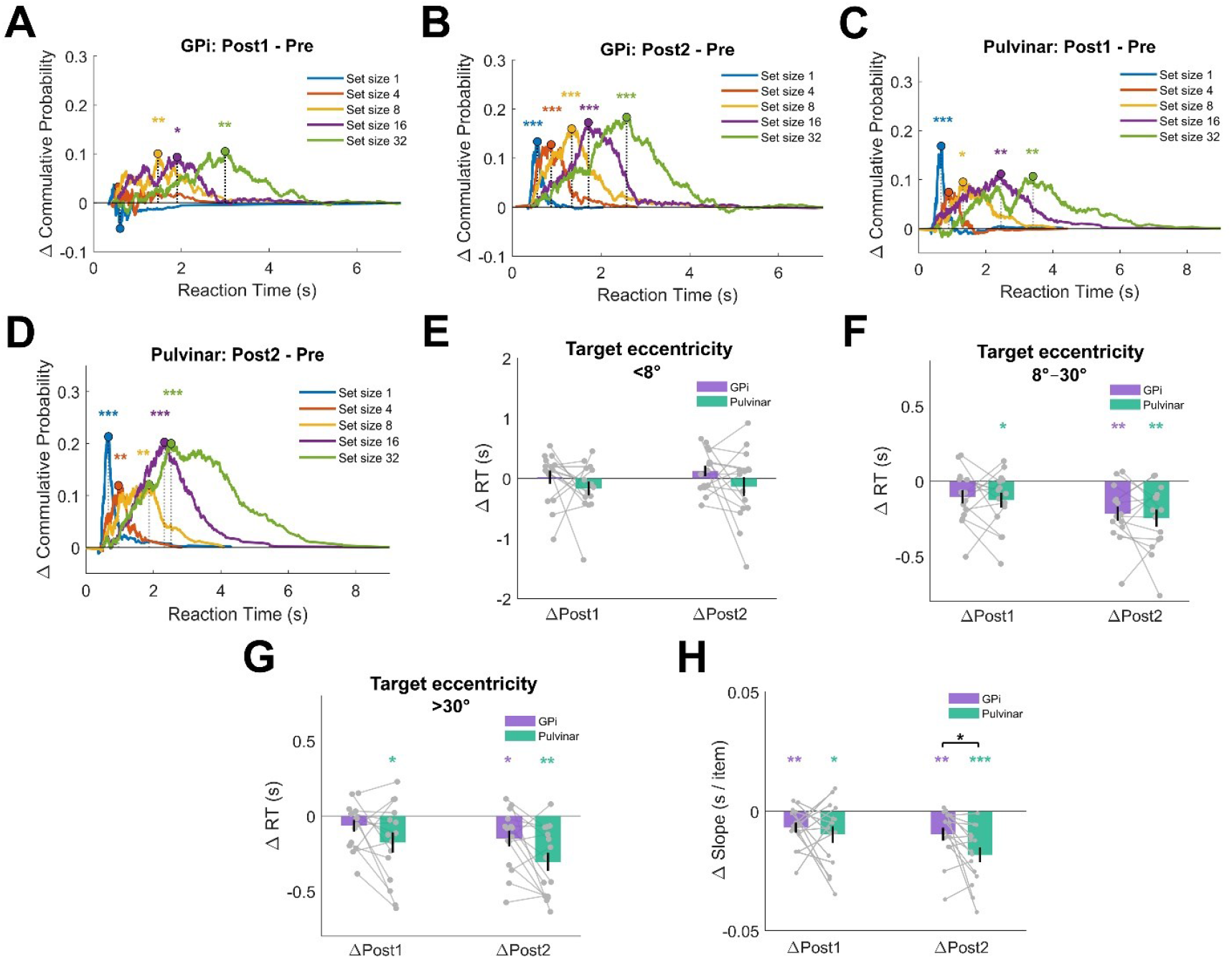
(A–D) Difference plots of cumulative RT distributions (Post–Pre) demonstrate systematic post-stimulation RT acceleration. Negative deflections indicate shifts of the underlying distributions toward shorter RTs. In GPi sessions, shifts were present at Post1 (A) and strongest at Post2 (B), particularly at higher set sizes (e.g., set size 32: ΔMedian = −0.478 s, KS D = 0.184, p = 2.6 × 10^−7^). Pulvinar stimulation also accelerated RTs, with significant changes already at Post1 (C) and the largest effects at Post2 (D), especially at high load (set size 32: ΔMedian = −0.468 s, KS D = 0.200, p = 2.7 × 10^−9^). The corresponding raw cumulative functions, which more intuitively illustrate the leftward shift in RTs, are shown in Fig. 3A (GPi) and Fig. 3B (Pulvinar), where target-present and target-absent trials are also displayed separately. (E-G) ΔRT (mean ± SEM) by target eccentricity. GPi stimulation produced significant speeding at intermediate eccentricities (8–30°) in Post2 (p = 0.0028), while pulvinar stimulation yielded stronger improvements at peripheral targets (>30°) in Post2 (p = 0.0019). A significant between-site difference emerged centrally (<8°) in Post1 (p = 0.03). (H) Search slope effects (s/item ± SEM). GPi slopes declined from 0.0586 to 0.0490 s/item at Post2 (Δ = −0.0096 ± 0.0105, p = 0.0065), while pulvinar slopes decreased more steeply, from 0.0680 to 0.0497 s/item (Δ = −0.0182 ± 0.0121, p = 8.5 × 10^−5^), with a significant between-site difference (p = 0.035). Significance stars: ***p < 0.001, **p < 0.01, *p < 0.05, n.s. = not significant (all values FDR-corrected). Bars and shaded error margins indicate SEM.

**Figure 3.**
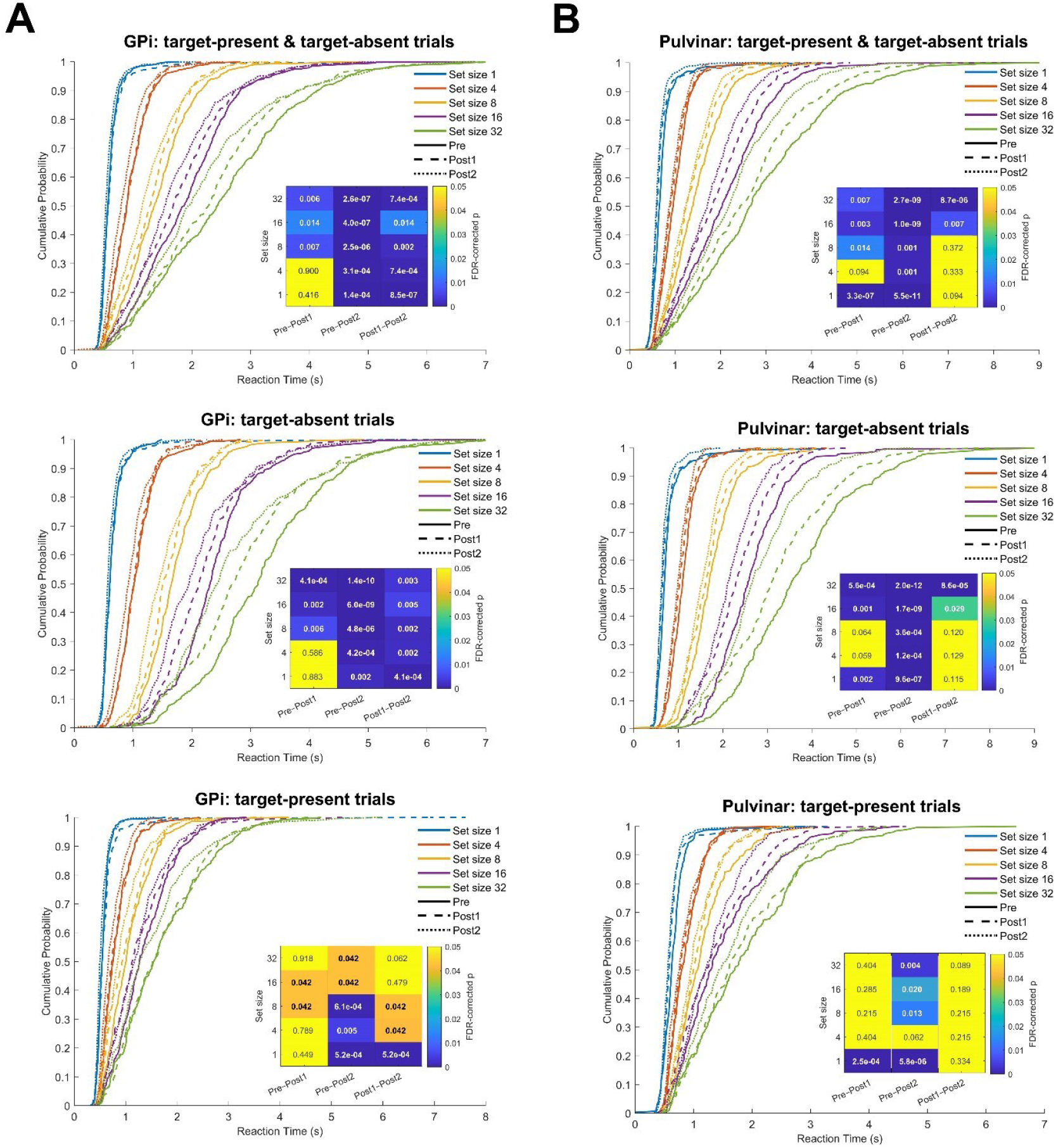
Cumulative distribution analyses of reaction times for GPi and pulvinar stimulation. (A) GPi. Empirical cumulative distribution functions (ECDFs) of reaction times across set sizes (1, 4, 8, 16, 32 items) for all trials combined (top), target-absent trials (middle), and target-present trials (bottom). Solid lines indicate Pre, dashed lines Post1, and dotted lines Post2 sessions. Insets report FDR-corrected p-values from Kolmogorov–Smirnov (KS) tests for Pre–Post1, Pre–Post2, and Post1–Post2 contrasts. (B) Pulvinar. ECDFs for all trials combined (top), target-absent (middle), and target-present (bottom) conditions. Insets show FDR-corrected KS p-values for each set size and session contrast.

We next examined how these improvements varied as a function of target eccentricity and trial type. Both GPi and pulvinar stimulation reduced average RTs at intermediate (8– 30°) and large (>30°) target eccentricities, but not for central targets (<8°, Fig. 2E-G). The larger benefits observed at more eccentric locations could reflect either a greater number of attentional shifts in the search for the target or simply that longer baseline RTs provided more opportunity for stimulation effects to accumulate. When trials were pooled, no reliable differences emerged between GPi and Pulvinar. However, when trials were separated by trial types (Fig. 4A–B), target-present trials (Fig. 4B) revealed a site-specific difference at peripheral locations (>30°), where pulvinar TUS produced stronger improvements than GPi TUS, suggesting that these areas may play distinct roles in supporting performance when targets appear in the far periphery.

**Figure 4.**
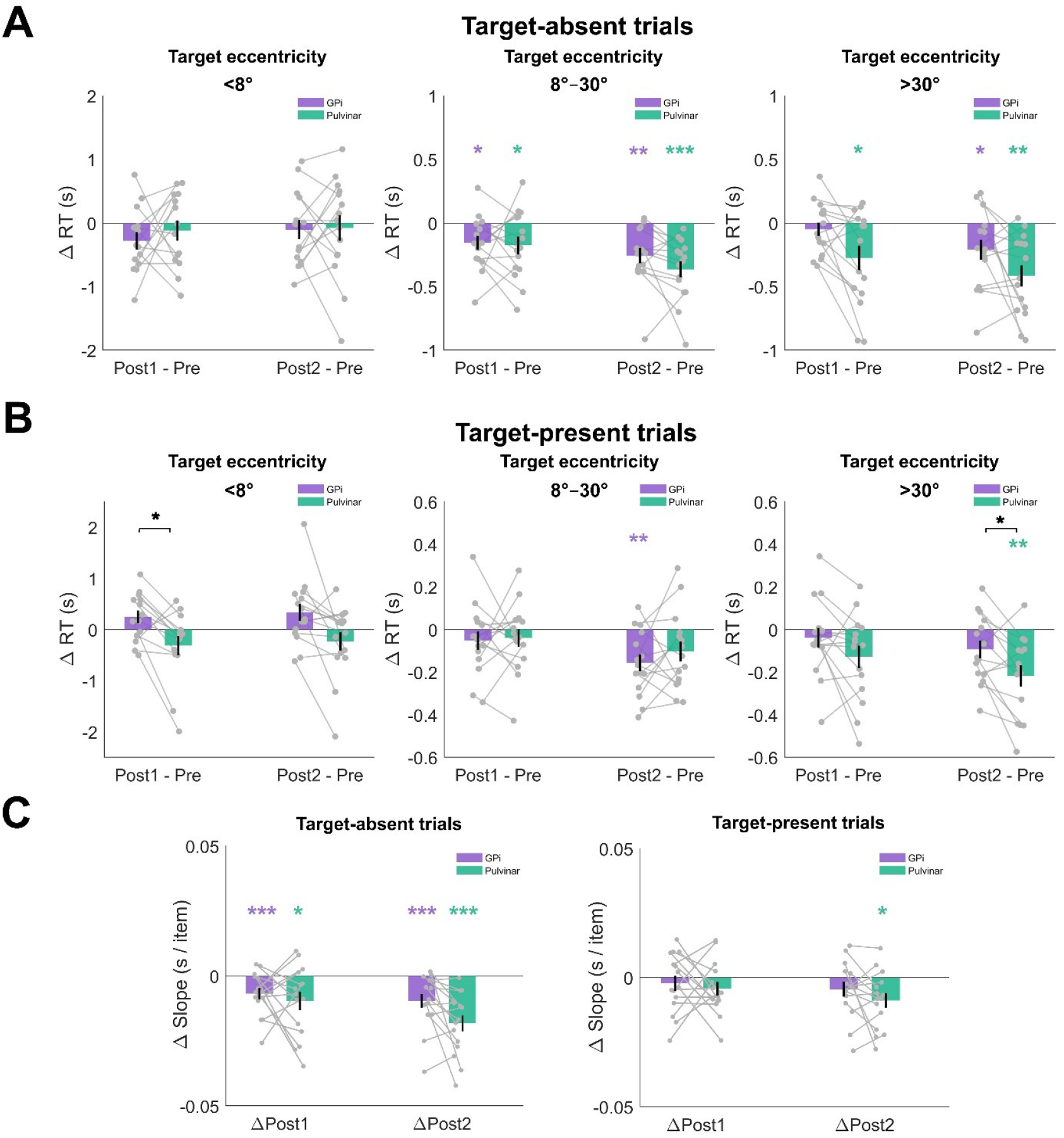
Eccentricity and search efficiency. (A) ΔRT by eccentricity for target-absent trials. Both GPi and pulvinar produced significant speeding at intermediate eccentricities and peripheral locations. (B) ΔRT by eccentricity for target-present trials. GPi showed facilitation at 8–30° (ΔPost2 p = 0.0078), while pulvinar selectively improved >30° (ΔPost2 p = 0.0071). A significant GPi–pulvinar difference was observed at <8° in Post1 (p = 0.03). (C) Target-absent trials (left plot): Both GPi and pulvinar showed significant slope reductions. GPi: ΔPost1 = −0.0111 ± 0.0074 s/item (p = 9.1 × 10^−5^), ΔPost2 = −0.0146 ± 0.0133 (p = 8.4 × 10^−4^). Pulvinar: ΔPost1 = −0.0150 ± 0.0204 (p = 0.013), ΔPost2 = −0.0272 ± 0.0187 (p = 1.2 × 10^−4^). Target-present trials (right plot): Pulvinar produced significant slope reductions at Post2 (Δ = −0.0089 ± 0.0110, p = 0.015), whereas GPi slope changes did not reach significance (all p > 0.5).

Search efficiency analyses further supported this dissociation. Slopes of the RT–set size function declined significantly after stimulation at both sites, reflecting increased processing efficiency per distractor, but the effect was stronger for pulvinar. GPi slopes decreased from 0.0586 s/item at baseline to 0.0490 s/item at Post2, corresponding to an increase in processing rate from 17.1 to 20.4 items/s (Δ = +3.3 items/s). Pulvinar slopes fell from 0.0680 to 0.0497 s/item, corresponding to an increase from 14.7 to 20.1 items/s (Δ = +5.4 items/s). This larger gain for pulvinar was reflected in a significant between-site difference (p = 0.035) (Fig. 2H). However, when trials were separated by trial types (Fig. 4C), only TUS of the pulvinar lead to increased search efficiency in the target-present trials. In the targe-absent trials, however, search efficiency was increased following stimulation of both areas (Fig. 4C).

## 4. Discussion

Our findings reveal that the human attentional system can be experimentally tuned at its deep subcortical sites. Within the pallido–thalamic network, both GPi and pulvinar stimulation accelerated reaction times at intermediate and peripheral target eccentricities. Importantly, a site-specific dissociation emerged in target-present trials, where pulvinar produced a more efficient search and stronger RT improvements than GPi at peripheral locations, suggesting that these structures may differentially support performance depending on spatial demands.

The pulvinar findings converge with a large body of work showing that this nucleus acts as an active attentional hub and contribute to selective filtering and prioritization of relevant inputs[18,20,22]. Human lesion and imaging studies demonstrate its role in suppressing distractors and amplifying relevant signals[20,21], and primate physiology reveals that pulvinar neurons synchronize cortical networks to bias perception[22]. Recent causal manipulations further support this vie; Boshra et al. (2025) showed that inducing burst firing in macaque pulvinar enhanced attentional performance [24]. Our results extend these findings to healthy humans, providing causal evidence that augmenting pulvinar function sharpens selective attention, particularly under high load and eccentricity. By contrast, GPi stimulation produced a more global speeding of responses, consistent with the basal ganglia’s role in regulating decision thresholds and urgency. Electrophysiological data indicate that pallidal neurons encode an urgency signal that accelerates commitment without representing specific stimulus features [30]. Anatomical studies also show GPi outputs reach prefrontal cortex[28], providing a substrate for influencing cognitive as well as motor functions. Our TUS data therefore support the idea that the GPi contributes to attentional performance by lowering global response thresholds, rather than improving spatial filtering. While our current study does not allow us to disentangle these mechanistic interpretations further, they can be directly tested in future experiments.

By directly and non-invasively stimulating these deep structures with focused ultrasound in healthy participants, we show that attentional control can be causally modulated without penetrating the brain. This work moves beyond correlational mapping and patient studies, offering, to our knowledge, the first within-subject demonstration that subcortical attention circuits can be differentially steered in the intact human brain. The results also resonate with our prior TUS work showing GPi, but not pulvinar, influences response inhibition[12], underscoring that even within a single anatomical network, distinct loops orchestrate separate facets of cognition. The dissociation between pulvinar and GPi also rules out nonspecific explanations: participants often reported drowsiness after stimulation, yet performance improved, opposite to what a generic arousal effect would predict. Moreover, the lack of significant difference in RTs between the first and second halves of trials within each block indicated that practice effects did not drive the speeding of responses, supporting the conclusion that TUS was responsible for the faster reaction times.

Methodologically, this study showcases the potential of TUS as a tool for cognitive neuroscience. Its millimeter-scale focality and ability to target structures several centimeters beneath the cortical surface allow manipulations previously restricted to invasive approaches. Mechanistically, TUS likely modulates neural activity through mechanosensitive ion channels and astrocytic signaling [32,33], producing offline effects that last beyond stimulation. While modeling minimized off-target engagement, neighboring structures (e.g., GPe, internal capsule, adjacent thalamic nuclei) cannot be fully excluded. Future improvements in beam steering and personalized dosing will refine targeting.

Beyond serving as a causal probe, TUS could be harnessed to augment cognitive functions. Enhancing pulvinar function may help mitigate distractibility in ADHD, while modulating GPi could support conditions with impaired response control such as Parkinson’s[34]. More broadly, if selective stimulation can enhance attention, it may also be possible to boost memory, mood, or other cognitive and affective domains by targeting their circuits. Once protocols are developed that yield longer-lasting effects, TUS could provide a noninvasive alternative to DBS, with both scientific and therapeutic impact.

## CRediT authorship contribution statement

**Hamidreza Ramezanpour**: Writing – review & editing, Writing – original draft, Visualization, Validation, Software, Resources, Project administration, Methodology, Investigation, Formal analysis, Data curation, Conceptualization.

**Ghazaleh Darmani**: Writing – review & editing, Visualization, Validation, Software, Project administration, Methodology, Investigation, Data curation, Conceptualization.

**Regina Annirood**: Writing – review & editing, Validation, Software, Project administration, Methodology, Investigation, Data curation.

**Can Sarica**: Writing – review & editing, Methodology, Investigation, Data curation.

**Jean-Francois Nankoo**: Writing – review & editing, Methodology, Investigation.

**Samuel Pichardo**: Writing – review & editing, Software, Resources, Methodology, Investigation.

**Jeffrey Schall**: Writing – review & editing, Validation, Resources, Conceptualization.

**Andres M. Lozano**: Writing – review & editing, Validation, Resources, Methodology.

**Robert Chen**: Writing – review & editing, Validation, Resources, Project administration, Methodology, Conceptualization.

## Acknowledgements

The study was funded by a Canadian Institutes of Health Research grants (FDN 154292, PJT 198046) and the Natural Science and Engineering Research Council grant (RGPIN-2020-04176) to R.C.

